# Spores-on-a-Chip: a new sample-loading strategy for multiplexed single-cell monitoring of spore germination within a microfluidic platform

**DOI:** 10.64898/2026.06.30.735560

**Authors:** Maria Portela, Claire E. Stanley

## Abstract

We demonstrate a new sample-loading strategy (“spot-loading”) for Spores-on-a-Chip microfluidic sensors that enables multiplexed experimentation. The previous limitation of one biological sample per device is overcome through controlled sample loading at an intermediate step of the chip fabrication process. As a proof-of-concept, we use dual spore chips to compare the germination behaviours of two spore strains in two distinct microenvironments.

## Introduction

Analytical tools for monitoring and studying microbial spores are important for a range of research applications, ranging from the food industry to healthcare. Due to their characteristic resistance to high temperatures, desiccation, radiation, and chemical attacks,^1^ bacterial spores have important applications in biotechnology,^2^ agriculture,^3^ and human health^4^. For example, spores can act as pesticides^5^, promote nutrient cycling in the soil,^3^ and contribute to bioremediation^6^ and carbon capture efforts.^7^ On the other hand, spores play key roles in the infection mechanism of bacterial pathogens such as *Clostridium difficile* and *Bacillus anthracis*.^4^ Spores of *C. difficile* can also resist common disinfectants, which is of particular concern in healthcare settings, where these resistant spores act as reservoirs and increase the risk of re-infection and disease transmission.^4^ In the food industry, some spore strains, such as *Bacillus cereus, Clostridium tyrobutyricum*, and *Alicyclobacillus acidoterrestris*, can survive pasteurisation, leading to food spoilage and increased disease outbreaks.^8, 9^ Spores may also play a role in antibiotic resistance, as they are better able to survive antibiotic exposure than fast-growing bacteria, and may germinate and resume growth once the treatment stops.^10^

Recent work has highlighted the role of single-cell heterogeneity in sporulation and germination processes. For example, Russel *et al*. used a “mother machine” microfluidic device to elucidate the role of stochastic noise in the signalling cascade which controls entry into sporulation in *Bacillus subtilis*, the model spore-forming bacterium.^11^ Mutlu and colleagues were able to immobilise spores on agar pads, and then image the life history of individual *B. subtilis* cells to find a “phenotypic memory” link between sporulation timing and spore revival.^12^ To better understand and control these processes, and thus develop improved decontamination and biocontrol strategies, there is a need for more versatile sensing platforms that allow single-cell monitoring of microbial sporulation and germination dynamics in a variety of application-relevant conditions.

We recently developed the 4-conditions microfluidic chemostat (4CMC) for the detection of microbial spore germination.^13^ This device uses laminar flow to create up to four separate regions with defined media conditions, across an array of observation chambers where spores can be trapped and monitored over time by microscopy. However, as samples are loaded via syringe injection, which results in a random distribution of spores across all observation chambers, the device is limited to investigating one biological sample at a time.

In the present work, we build upon the 4CMC device, expanding its capabilities for use in multiplexed Spores-on-a-Chip experiments. Instead of relying on syringe loading, we used a new sample-loading strategy – “spot-loading” – that exploits spores’ characteristic resistance to desiccation. Spores were loaded in a controlled manner, at an intermediate step of the device fabrication process. The resulting multiplexed devices were benchmarked against devices loaded conventionally, and tested in a proof-of-concept experiment that allowed direct detection of germination events for two spore samples under two different media conditions, at the single-cell level.

## Results and discussion

To manufacture multiplexed Spores-on-a-Chip devices, we adapted the method conventionally used for making poly-(dimethylsiloxane) (PDMS) microfluidic devices. First, we cast PDMS slabs from a master mould containing the device design, an adapted, 2-inlet version of the 4CMC device^13^ (Figure 1A). To fabricate devices using the conventional method, we bonded PDMS slabs directly to clean glass coverslips using a plasma oven, and then loaded samples via syringe injection (Figure 1B), using about 200 µL of a spore suspension to account for sample losses in the syringe and tubing. In contrast, when fabricating chips using the spot-loading technique, we also plasma-treated a clean glass coverslip, but then applied a methanol treatment, which was found to increase surface hydrophobicity while still allowing good bonding to the plasma-activated PDMS. The surface contact angle of water on the glass increased from under 5° after plasma treatment, to 22.4° ± 0.4° after methanol treatment (n = 8 coverslips, with 5-7 droplets each). This allowed the deposition of multiple 1 µL droplets containing a spore suspension onto the glass surface, with a well-defined shape and in a controlled position. After the spore droplets dried out completely, the PDMS slab was plasma-activated and placed on top of the spotted spores, with the array of observation chambers aligned to the spore suspension spots, to seal the microfluidic device (Figure 1C). Because the area of the spore spots was much larger than the area of the chambers, we found that some spores were trapped under the pillars that flank each observation chamber. This limited the maximum loading concentration for spot-loaded devices: while most pillars bonded correctly with an optical density at 600 nm (OD600) of 0.5 or lower, higher concentrations led to poor-quality devices, as spore clumps prevented contact between the PDMS slab and the glass surface, which is required for good bonding. Further work to match spore spot and chamber areas could overcome this limitation, for example by designing larger chambers, using precision dispensing systems to decrease spot size, or using emerging surface patterning techniques to better control spotting geometry.^14^ We then characterised the spot-loading strategy, comparing it with conventional loading. While in both cases a higher loading concentration led to a higher median number of spores per chamber, conventional loading required a concentration one order of magnitude larger to achieve similar numbers of spores per chamber as spot-loading (Figure 1D), likely due to sample losses in the syringe and tubing with conventional loading. As the spot-loading strategy leads to a broader distribution of spores per chamber, it may be better suited to studies where having a variety of starting spore numbers is important, such as investigations on spore-spore interactions, or where only small amounts of sample material are available. We also noted that the proportion of chambers with at least one spore decreased sharply at lower loading concentrations, particularly for conventionally loaded devices (Figure 1E), which decreases the statistical power of experiments. In contrast, for spot-loaded devices at an OD600 of 0.05 or above, nearly all chambers within the spot area (about 35% of the total observation area) had at least one spore. This suggests that increasing the size or number of spore spots could increase the chamber occupancy rate to nearly 100%, maximising the amount of data that can be collected with a single device.

**Fig. 1:**
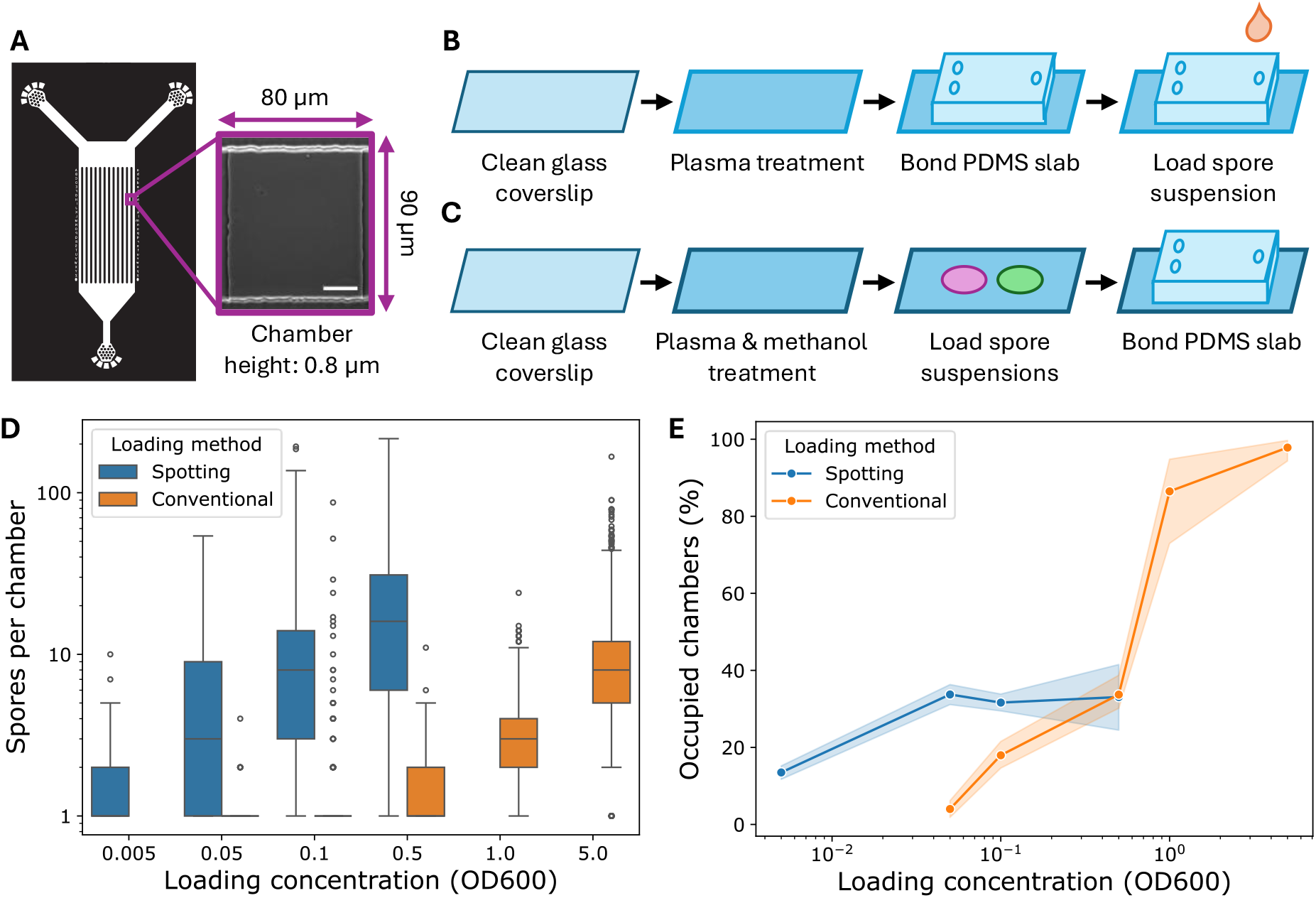
Characterising the spot-loading strategy. A) Design of the microfluidic device used in this study, showing the two inlets (top), the outlet (bottom), and the array of observation channels perfused by 12 parallel supply channels. Laminar flow in the devices prevents mixing of the media perfused through each inlet. Detail: phase-contrast micrograph showing an example observation chamber. The height of the supply channels is 10 µm, whereas the observation chambers are 0.8 µm high. The scale bar is 20 µm. B) Schematic of the conventional device fabrication and loading protocol. C) Schematic of the spot-loading strategy. D) Comparison of the number of mRFP1-tagged *B. subtilis* spores loaded in each observation chamber, by loading concentration and method (n = 4-6). Only chambers with at least one spore are included. The differences between all groups are significant (Mann-Whitney two-sided test, p < 0.001), except for conventional devices loaded at OD600 0.05 and 0.1. E) Comparison of the proportion of chambers in each device occupied with at least one spore, by loading concentration and method. The shaded area represents the 95% confidence interval (n = 4-6).

After characterising our newly developed spot-loading strategy, we investigated whether it affects the viability and germination behaviour of spores. To do this, we conducted germination experiments using devices spot-loaded with spores of *B. subtilis*, the model spore-former bacteria. We compared the germination and outgrowth of these spores in a nutrient-rich broth (NB), and a nutrient-poor media (SM), which had no added amino acids. In our experiments, spores switched from a phase-bright to a phase-dark appearance in both conditions, characteristic of the first stages of spore revival, and in nutrient-rich media cells also increased in size, a sign of outgrowth, one of the final stages of spore revival (Figure 2A and Movies S1-S2).^15^ Using the PySpore^16^ software, we quantified germination parameters for the individual spores tracked in our experiments. Our results show spore traces with sharp drops in pixel intensity over time, indicating germination events, with nutrient-rich media leading to a larger average decrease in intensity (Figure 2B). This corresponds to a higher germination rate (Figure 2C), and a larger proportion of germinated spores after 1.5h of media exposure (Figure 2D) in nutrient-rich media. These results are consistent with previous findings that certain small molecules, such as amino acids and sugars, bind germination receptors at the spore surface to trigger germination.^17^ These receptors have different specificities and often cooperate, so a richer media is likely able to trigger more receptors, leading to faster and more frequent germination.^1, 18^ Some germination may still be expected in nutrient-poor media, as glucose is recognised by germination receptors in *B. subtilis*, although the exact mechanism is not fully understood.^17, 19^ Our data also shows a significant increase in cell area over time for the spores that germinated in nutrient-rich media (Figure 2E), with cell area growing by nearly 40% in the 100 minutes after germination, compared to an 8% increase in cell area for the nutrient-poor media (Figure 2F). This indicates that cell outgrowth occurred only in the nutrient-rich media, in line with previous findings that extrinsic amino acids are required for vegetative growth in newly germinated spores.^20^ Therefore, our findings match the expected response under the conditions tested, with spore germination faster and more frequent in nutrient-rich media, and outgrowth only present when there are amino acids present to support vegetative cell growth.

**Figure 2:**
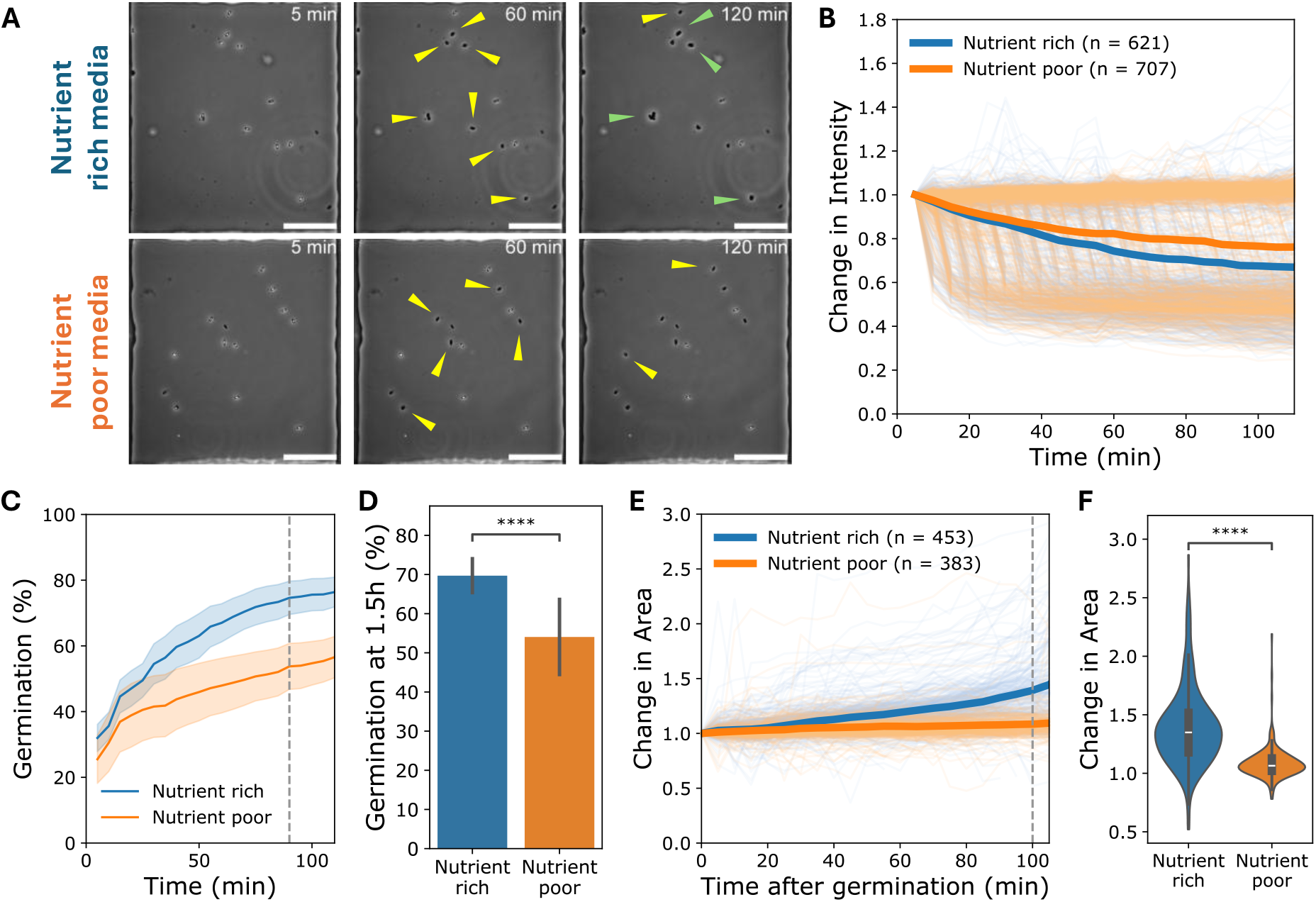
Germination in spot-loaded devices. A) Phase-contrast timelapses of two representative chambers in a device spot-loaded with an OD600 0.05 spore suspension of mRFP1-tagged *B. subtilis*. The scale bar is 20 µm. Spores are phase-bright, germinated cells are phase-dark. Yellow arrows show germinated cells, and green arrows shows cells that have increased in size. B) Change in pixel intensity of individual spores tracked throughout germination experiments. Higher values represent phase-bright spores. Thin lines represent individual cells, and thick lines show the mean for each media condition (data from 6 devices). C-D) Percentage of germinated cells in each observation chamber over time (C) and at time t = 100 min (D), across 6 devices. Shaded area and error bars represent the standard error (n = 6). Two-tailed proportions Z-test (n = 621 or 707 individual cells, for rich/poor media). E-F) Change in individual cell area after spore germination over time (E) and at time t = 100 mins after spore germination (F), across 6 devices. Time t = 0 is taken as the germination time point for each individual cell. Two-tailed independent t-test (n = 236 or 207 individual cells, for rich/poor media).

Having developed and validated a spot-loading strategy to fabricate Spores-on-a-Chip devices loaded with more than one biological sample (Figure 3A and Movies S3-S6), we then investigated its potential for multiplexed germination assays. To do this, we set-up a proof-of-concept experiment with spores from two different bacterial species, *B. subtilis* and *Ammoniphilus oxalaticus*, and two different chemical stimuli, L-alanine or potassium oxalate, so that we could investigate the responsiveness of each type of spore to each chemical stimulus (Figure 3B). While *B. subtilis* has long been known to germinate in the presence of L-alanine,^21^ spores of *A. oxalaticus* require oxalate to germinate.^22^ Our experiment corroborated these previous findings: germination occurred for *A. oxalaticus* exposed to potassium oxalate, and for *B. subtilis* exposed to L-alanine; but was otherwise very low or absent (Figure 3C). We did not observe mixing of the spore samples, which in this experiment would have been evident due to the significant difference in size between the two spore strains tested. After 2 h, about 35% of the *A. oxalaticus* spores exposed to oxalate had germinated, while all the spores exposed to L-alanine remained dormant. In contrast, just over 25% of the *B. subtilis* spores germinated in response to L-alanine, with only about 4% germinating in response to oxalate-containing media (Figure 3D). One factor we did not examine is whether exudates from germinating cells in chambers closer to the inlets are able to diffuse into other chambers and affect the spores there. Future studies using computational fluid dynamics to investigate how parameters such as exudate concentration and flow rate through the supply channels impact the diffusion rate of exudates into downstream chambers may provide useful insights. However, the potential confounding effect of these exudates can be mitigated with careful experimental design choices, for example by adding repeats where the order of spore spots is reversed, and controlling for position along the supply channel during data analysis.

**Figure 3:**
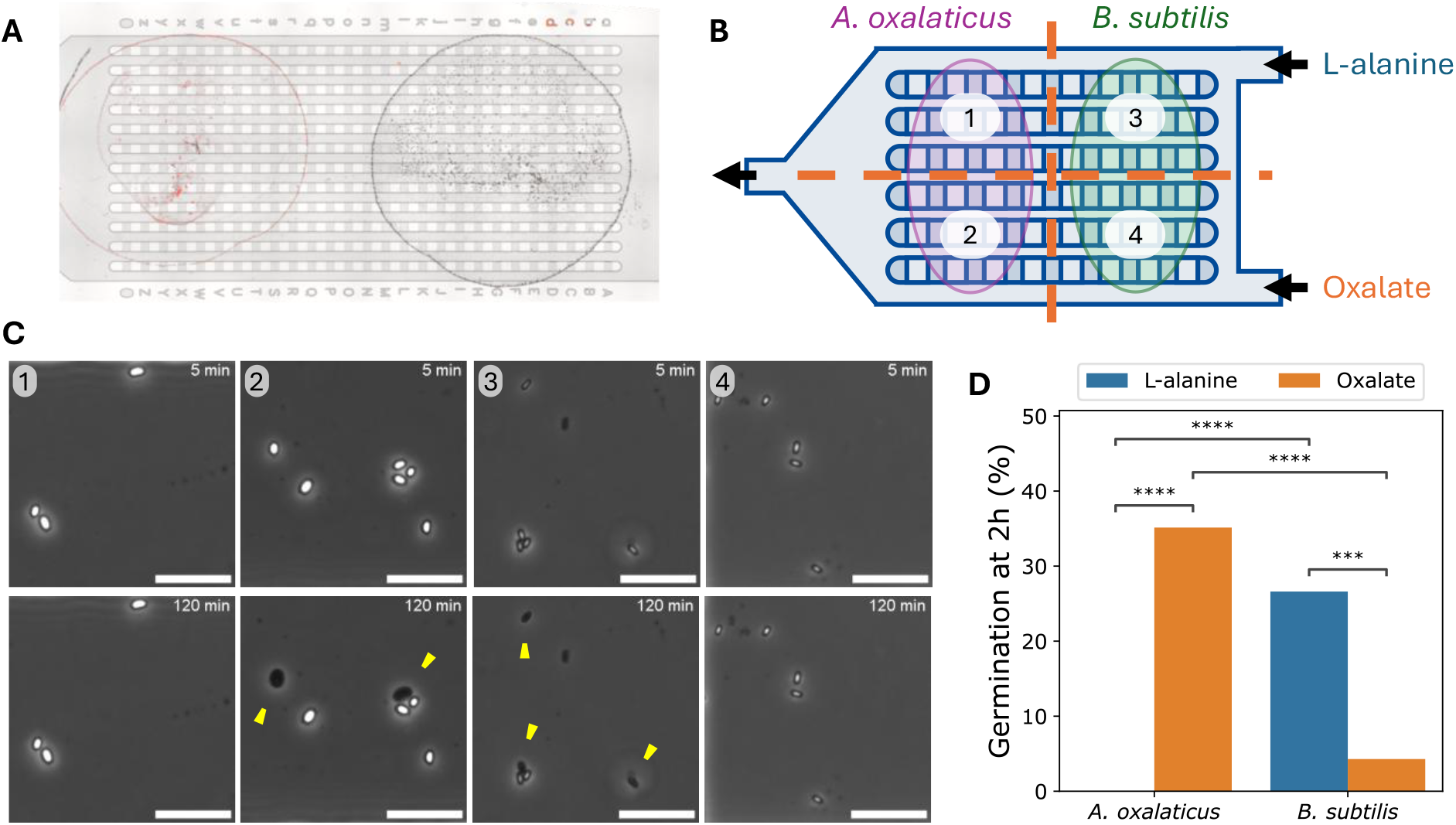
Multiplexed Spore-on-a-Chip experiments. A) Representative whole-device image of a dual Spores-on-a-Chip device, with RFP-tagged (left) or wild-type (right) *B. subtilis* 168. B) Schematic showing a multiplexed Spores-on-a-chip experiment, where spot-loaded samples of *A. oxalaticus* HJ and mRFP1-tagged *B. subtilis* spores, at a loading concentration of OD600 0.5, were continuously perfused with either 10 mM L-alanine in SM media or 0.65 g/L potassium oxalate in Schlegel AB media. This setup creates four combinations of two spore strains and two media conditions, numbered 1 to 4. C) Representative germination timelapses from zones 1 to 4 (see panel B). The scale bar is 10 µm. Yellow arrows highlight cells that germinated over the course of the experiment. D) Percentage of germinated cells in each observation chamber after 2h. Statistical testing: pairwise two-tailed proportions Z-tests (n = 59, 74, 109 and 70 individual cells, respectively for groups 1-4, data from 1 device).

## Conclusions

Taken together, our results show that the spot-loading strategy, besides enabling multiplexed experiments, also reduces the amount of biological material needed per experiment by roughly 2000-fold, is particularly well-suited to detection problems examining the effect of spore-to-spore proximity or where spore starting material is low or difficult to acquire, and increases the amount of data that can be obtained with a single device, without significantly impairing the viability or the normal germination biology of spore samples. We also show how the spot-loading strategy, if used with appropriate experimental design considerations, enables multiplexed germination sensing experiments that can directly compare how different spore samples respond to a set of defined media conditions, at the single-cell level, and within the same device.

Although we focussed predominantly on demonstrating a proof-of-concept application with two samples, the spot-loading strategy overcomes the fundamental limitation of one sample per device imposed by conventional syringe loading. Future experiments using a larger number of small spore spots could achieve higher throughputs, with the number of samples in a single device now limited by the number of spore spots that can be deposited on a glass coverslip (which in turn is determined by the minimal size of suspension droplets that it is possible to produce), and how well these spots can be aligned to a microfluidic chamber. Future studies should be able extend this technique to fungal spores. Besides multiplexing, this technique also enables co-culture studies: by depositing two overlapping spore spots, it is possible to quantify the response of each individual spore strain, as well as the resulting co-culture, in a single device. The spot-loading strategy therefore broadens the potential applications of Spores-on-a-Chip sensor devices, making it a more versatile screening and diagnostic platform for spore germination and monitoring studies.

## Experimental

### Strains

*B. subtilis* 168 (ATCC 23857), *B. subtilis* 168 expressing mRFP1, and the environmental isolate *A. oxalaticus* HJ.^22^

### Media

Nutrient broth (NB, Oxoid CM0001); synthetic minimal (SM): 1 mM D-glucose, 1 mM D-fructose and 10 mM KCl.^13^ Schlegel AB: ammonium, essential salts and trace elements.^22^

### Microfluidic device fabrication

PDMS (Dow, Sylgard 184) was prepared using a 10:1 ratio of base to curing agent, and bonded using a Zepto plasma cleaner (Diener Electronic; 1 min treatment, 0.75 mbar vacuum pressure, 50% power) onto clean borosilicate glass coverslips (VWR). Methanol treatment for spot-loading: 37 kHz sonication for 5 mins (Elmasonic S30).

### Live-cell imaging

Phase-contrast imaging with a Nikon Eclipse Ti2 microscope, with a stage incubator at 37°C (or 30°C for *A. oxalaticus*). Imaging started 5 mins after the start of media perfusion at 5 µL/min via a syringe pump (AL-1600).

Statistical annotations: ***: p < 0.001 and ****: p < 0.0001.

Full details are provided in the supplementary information.

## Author contributions

**Maria Portela:** Conceptualisation, Methodology, In estigation, Software, Visualisation, Formal analysis, Writing original draft; **Claire**. **Stanley:** Conceptualisation, Funding acquisition, Super ision, Writing – re iewing editing.

## Data availability

Raw data, processed files, and analysis scripts available on Zenodo at: https://doi.org/10.5281/zenodo.20734901.^23^

## Acknowledgements

We acknowledge financial support from the Department of Bioengineering and a President’s Ph Scholarship (Imperial College ondon, UK), as well as Prof. Tom Ellis (Imperial College London, UK) and Prof. Pilar Junier (University of Neuchâtel, Switzerland) for providing bacterial strains used in this study.

