## Supplementary Information for "Spores-on-a-Chip: a new sample-loading strategy for multiplexed single-cell monitoring of spore germination within a microfluidic platform"

### Supplementary Movies

#### Supplementary Movie 1

**Description: Responsiveness of spot-loaded mRFP1-tagged *B. subtilis* spores under nutrient-rich conditions.** Phase-contrast timelapses of a representative observation chamber in a Spores-on-a-Chip device spot-loaded with an OD600 0.05 spore suspension of mRFP1-tagged *B. subtilis*, and perfused with nutrient broth (NB) at a rate of 5 µm/min. Image acquisition started 5 minutes after the start of media perfusion, and images were acquired at 5-minute intervals over 2 hours. The movie shows spore germination (phase-bright to phase-dark switch) and outgrowth. The scale bar is 20 µm.

#### Supplementary Movie 2

**Description: Responsiveness of spot-loaded mRFP1-tagged *B. subtilis* spores under nutrient-poor conditions.** Phase-contrast timelapses of a representative observation chamber in a Spores-on-a-Chip device spot-loaded with an OD600 0.05 spore suspension of mRFP1-tagged *B. subtilis*, and perfused with synthetic minimal media (SM) at a rate of 5 µm/min. Image acquisition started 5 minutes after the start of media perfusion, and images were acquired at 5-minute intervals over 2 hours. The movie shows spore germination (phase-bright to phase-dark switch) but no evidence of spore outgrowth post-germination. The scale bar is 20 µm.

#### Supplementary Movie 3

**Description: Responsiveness of spot-loaded *A. oxalaticus* spores exposed to L-alanine.** Phase-contrast timelapses of a representative observation chamber in a multiplexed Spores-on-a-Chip device spot-loaded with an OD600 0.05 spore suspension of *A. oxalaticus*, and perfused with 10 mM L-alanine at a rate of 5 µm/min. Image acquisition started 5 minutes after the start of media perfusion, and images were acquired at 5-minute intervals over 2 hours. The movie shows no evidence of spore germination. The scale bar is 20 µm.

#### Supplementary Movie 4

**Description: Responsiveness of spot-loaded *A. oxalaticus* spores exposed to potassium oxalate.** Phase-contrast timelapses of a representative observation chamber in a multiplexed Spores-on-a-Chip device spot-loaded with an OD600 0.05 spore suspension of *A. oxalaticus*, and perfused with 0.65 g/L mM potassium oxalate at a rate of 5 µm/min. Image acquisition started 5 minutes after the start of media perfusion, and images were acquired at 5-minute intervals over 2 hours. The movie shows spore germination. The scale bar is 20 µm.

**Supplementary Movie 5**

**Description: Responsiveness of spot-loaded *B. subtilis* spores exposed to L-alanine.** Phase-contrast timelapses of a representative observation chamber in a multiplexed Spores-on-a-Chip device spot-loaded with an OD600 0.05 spore suspension of *B. subtilis*, and perfused with 10 mM L-alanine at a rate of 5 µm/min. Image acquisition started 5 minutes after the start of media perfusion, and images were acquired at 5-minute intervals over 2 hours. The movie shows spore germination. The scale bar is 20 µm.

**Supplementary Movie 6**

**Description: Responsiveness of spot-loaded *B. subtilis* spores exposed to potassium oxalate. Phase**-contrast timelapses of a representative observation chamber in a multiplexed Spores-on-a-Chip device spot-loaded with an OD600 0.05 spore suspension of *B. subtilis*, and perfused with 0.65 g/L mM potassium oxalate at a rate of 5 µm/min. Image acquisition started 5 minutes after the start of media perfusion, and images were acquired at 5-minute intervals over 2 hours. The movie shows no evidence of spore germination. The scale bar is 20 µm.

### Supplementary Methods

#### Reagents

| **Reagent** | **Supplier** | **Catalogue #** |
|---|---|---|
| Nutrient broth (NB) | Oxoid | CM0001 |
| Nutrient agar (NA) | Oxoid | CM0003 |
| D-fructose | Millipore | 47740 |
| D-glucose | Sigma-Aldrich | G7528 |
| Potassium chloride (KCl) | Fisher BioReagents | BP366500 |
| Sodium phosphate dibasic ($Na_2HPO_4$) | Sigma-Aldrich | S5136 |
| Monopotassium phosphate ($KH_2PO_4$) | Sigma-Aldrich | P5655 |
| Ammonium chloride ($NH_4Cl$) | Sigma-Aldrich | A4514 |
| Magnesium sulphate heptahydrate ($MgSO_4.7H_2O$) | Sigma-Aldrich | M2773 |
| Ammonium iron(III) sulphate dodecahydrate ($Fe(NH_4)(SO_4)_2.12H_2O$) | Sigma-Aldrich | F3629 |
| Calcium chloride dihydrate ($CaCl_2.2H_2O$) | Sigma-Aldrich | C3306 |
| Zinc sulfate heptahydrate ($ZnSO_4.7H_2O$) | Sigma-Aldrich | Z0251 |
| Manganese(II) chloride tetrahydrate ($MnCl_{2.4}H_2O$) | Sigma-Aldrich | 63535 |
| Boric acid ($H_3BO_3$) | Sigma-Aldrich | B6768 |
| Cobalt(II) chloride hexahydrate ($CoCl_2.6H_2O$) | Sigma-Aldrich | C2911 |
| Copper(II) chloride dihydrate ($CuCl_2.2H_2O$) | Sigma-Aldrich | C3279 |
| Nickel(II) chloride hexahydrate ($NiCl_2.6\ H_2O$) | Sigma-Aldrich | 223387 |
| Sodim molybdate dihydrate($Na_2MoO_4.2\ H_2O$) | Sigma-Aldrich | M1651 |
| Agar | Sigma-Aldrich | A1296 |
| Calcium oxalate | Sigma-Aldrich | 289841 |
| Potassium oxalate | Sigma-Aldrich | P0963 |
| L-alanine | Sigma-Aldrich | A7469 |
| HistoDenz | Sigma-Aldrich | D2158 |
| Fluorescein sodium salt | Sigma-Aldrich | 46960 |

| | | |
|---|---|---|
| SU-8 2002 | Kayaku Advanced Materials | 2002 |
| SU-8 2010 | Kayaku Advanced Materials | 2010 |
| Poly-dimethylsiloxane (PDMS) | Dow | Sylgard 184 |
| Sodium hydroxide (NaOH) | Thermo Fisher | BP359 |
| Ethanol | VWR | 20821.330 |
| Methanol (≥99.8%), ACS Analytical Reagent | VWR | 20847.307 |
| Isopropanol (≥99.7%), ACS Analytical Reagent | VWR | 20842.323 |
| Acetone (≥99%), Technical | VWR | 20063.365 |

**Bacterial strains and spore purification**

Two strains *Bacillus subtilis* were used in this study: a wild-type *B. subtilis* 168 strain (ATCC 23857), and a mRFP1-tagged *B. subtilis* strain (vegP/spoVK promoter on a pNW33N backbone, derived from a B. subtilis 168 parent strain), constructed and provided by Prof. Tom Ellis' lab (Imperial College London, UK). Plate cultures of the environmental isolate *Ammoniphilus oxalaticus* HJ[1] were provided by Prof. Pilar Junier's lab (University of Neuchâtel, Switzerland).

To obtain *B. subtilis* spores, plate cultures in nutrient agar supplemented with 5 g/L chloramphenicol were incubated at 30° C for 3-4 weeks. Cultures were then heat-treated at 70 °C for 30 minutes to deactivate remaining vegetative cells, and spores were harvested and washed three times (4000 rpm centrifugation for 5 min) with autoclaved Milli-Q water. Spores were further purified using a density gradient protocol: the pellets were resuspended in 20% (w/v) HistoDenz (Sigma), gently pipetted onto a 2-layer gradient of HistoDenz (40% layered on 50% w/v) and centrifuged at 4000 rpm for 90 minutes. The resulting pellet was washed a further 3 times, resuspended in autoclaved Milli-Q water and stored at 4 °C until use.

To obtain spores of *A. oxalaticus* HJ, cells were first cultured on bilayered Schlegel AB plates supplemented with 4 g/L calcium oxalate. Plate cultures were incubated at room temperature for eight weeks. Spores were then harvested in 2 mL autoclaved Milli-Q water, washed (4000 rpm centrifugation for 10 min) five times in autoclaved Milli-Q water, and used immediately. HistoDenz centrifugation was not used to avoid sample loss.

**Media recipes**

Nutrient Broth (NB) and Nutrient Agar (NA) were prepared according to supplier's instructions and sterilised by autoclaving. Synthetic minimal media (SM) was prepared with 1 mM D-fructose, 1 mM D-glucose and 10 mM potassium chloride in autoclaved Milli-Q water, and then sterilised with 0.22 µm filters (Minisart NML, Sartorius).

Liquid Schlegel AB media consisted of 67% Schlegel A and 1% Schlegel B in autoclaved Milli-Q water. Schlegel A is prepared with 9 g/L sodium phosphate dibasic, 1.5 g/L monopotassium phosphate, 1 g/L ammonium chloride, 0.2 g/L magnesium sulphate, and 1 mL/L 10x trace elements solution. Schlegel B is prepared with 0.92 g/L ammonium iron sulphate and 1.12 g/L calcium chloride. The 10x trace elements solution was prepared with 1 g/L zinc sulphate, 0.3 g/L manganese chloride, 3.0 g/L boric acid, 2.0 g/L cobalt chloride, 0.1 g/L copper chloride, 0.2 g/L nickel chloride, and 0.3 g/L sodium molybdate. Bi-layered Schlegel AB plates were prepared with a layer of Schlegel AB with 4g/L calcium oxalate and 15 g/L technical agar, on top of a layer of Schlegel AB with 15 g/L agar with no added calcium oxalate, at a 1:3 ratio.

For germination experiments with *B. subtilis*, we used NB, Synthetic Minimal (SM) media, and SM supplemented with 10 mM L-alanine. For germination experiments with *A. oxalaticus*, we used Schlegel AB, with or without 0.65 g/L potassium oxalate supplementation. All solutions were sterilised using 0.2 µm filters.

**Microfluidic device fabrication**

Microfluidic chemostat devices were fabricated as previously described[2], using a version of the 4CMC device design with two inlets. Briefly, to create a master mould, a mask forming a pattern of observation chambers was fabricated in a 4-inch silicon wafer (Si-Mat Silicon Materials) by photolithography with SU-8 2002 photoresist. The un-masked area was then etched to a height of 0.76 ± 0.01 µm using $C_4F_8/SF_6$ plasma in an ICP chamber (Oxford Instruments, PlasmaPro 100 Cobra). Finally, 10 µm features forming inlets, supply channels, and the outlet, were fabricated by photolithography using SU-8 2010. Feature heights were verified using a Veeco Dektak 6M stylus profilometer.

For both types of devices, poly(dimethylsiloxane) (PDMS) slabs (prepared with a 10:1 ratio of base to curing agent) were then cast from this master mould and cured at 70 °C overnight, then diced, and holes for the inlets and outlets were punched with a precision cutter (Ø 1.02 mm; Syneo, USA). The slabs were then washed sequentially by 3 mins sonication in 0.5 M sodium hydroxide, 70% ethanol, and Milli-Q water (37 kHz, Elmasonic S30, Elma Schmidbauer GmbH), and dried at 70 °C overnight. Glass coverslips (VWR, borosilicate glass) were washed sequentially, by sonication for 5 mins, in acetone, isopropanol (IPA) and twice in Milli-Q water, and dried overnight at 70 °C.

To fabricate devices for conventional loading, clean glass coverslips and PDMS slabs were plasma-treated for 1 min in a Zepto plasma cleaner (Diener Electronic; vacuum pressure 0.75 mbar, power 50%), and then bound together. Samples were manually loaded into the device inlet by syringe injection via the outlet, using about 200 µL of spore suspension per device.

To fabricate spot-loaded devices, clean glass coverslips were plasma-treated for 1 min in the Zepto plasma cleaner, sonicated in methanol for 5 mins and then placed in a hot plate at 110 °C to dry. Coverslips were left to cool down for at least 10 mins before spore suspensions were deposited on the surface, in 1 µL droplets. For dual chips, droplets were deposited 2-3 mm apart, so that spots are separate but within the device observation zone. Droplets were air-dried completely for at least 10 mins and then a PDMS slab was plasma-treated for 1 min and bound on top of the spore spots, sealing the device. The devices were placed in a hot plate at 50 °C for 5 mins, to improve bonding quality, before further use.

Flow patterns within both types of microfluidic devices were characterised using 0% and 0.25% fluorescein solutions in Milli-Q water, and we observed that a laminar flow pattern developed correctly in both cases, with no fluid mixing.

The contact angles of Milli-Q water on treated glass coverslips were measured using a contact angle goniometer (Ossila L2004A1). A 10 µL droplet was deposited on the treated surface, and then we used Ossila's Contact Angle software (v3.0) for measurements. Bonding quality was tested with peel tests to check if the PDMS slab could be separated from the coverslip, both before and after filling the device with Milli-Q water. We also performed leak tests, manually flushing devices with a with Milli-Q water and inspecting for leaks.

**Live-cell imaging**

Devices were imaged with a Nikon Eclipse Ti2 microscope equipped with a camera (DS-Qi2, Nikon), and a TRITC-B (EX 543/22, DM 562, EM 593/40) filter cube, using the NIS Elements Advanced Research (v5.41.01) software from Nikon. We used a Plan Fluor 10x/0.3 or a Plan Fluor 20x/0.45 objective lens with the Ph1 phase contrast module for whole-device images, and an oil-immersion Plan Fluor 100x/1.3 objective (Nikon) with the Ph3 phase contrast module for timelapses of individual observation chambers. Acquired .nd2 files were converted to .tiff files using the NIS Elements Viewer (v5.21.00) software from Nikon.

**Data processing**

To quantify the distribution of spores in chambers, we developed a custom macro in Jython, using Fiji (v2.14.0, with ImageJ v1.54 and Java v1.8.0_322)[3]. Spores were detected by their red fluorescence. The macro asks for user input to determine a fluorescence threshold and define the observation area, and then identifies individual chambers and quantifies the number of fluorescent particles inside each chamber. A watershed transformation is applied to improve segmentation accuracy of closely spaced spores. Large area outliers, corresponding to spore clumps, were excluded from further analysis.

To analyse spore germination, we first used a custom Jython macro in Fiji to pre-process all videos. This pre-processing step converted images to an 8-bit format required for subsequent analysis steps and applied the StackReg[4] drift correction algorithm using a rigid body transformation. Manual drift correction was applied in a few cases where needed. Videos were then manually cropped to exclude chamber edges, which would have introduced segmentation artifacts in downstream analysis. Areas with illumination artifacts that didn't contain spores were also cropped out.

We then used the PySpore[5] software to segment and track individual cells, and extract germination metrics over time. All videos were manually checked for accurate segmentation at representative frames. Videos with frames significantly out-of-focus, or with significant oscillations in illumination, were excluded. For 8 videos, a single outlier frame with significant focus or illumination problems was excluded and replaced by the previous frame. We found that PySpore didn't perform very well on *A. oxalaticus* timelapses, often losing track of spores over time. For these experiments (Figure 3), germination counts were established manually: the number of spores and germinated cells at the 5 min, 2h and 12h timepoints, was counted and recorded for each timelapse video.

**Data analysis and visualisation**

Data exported from the ImageJ macros was further processed with Python (v3.11.9) in custom Jupyter Notebooks, using the Pandas (v2.2.3)[6], Numpy (v2.0.1)[7] and Scipy (v1.15.1)[8] packages. Plots and statistics were produced with Matplotlib (v3.10.1),[9] Seaborn (v0.13.2),[10] Statsmodels (v0.14.4)[11] and Statannotations (0.7.1).[12]
